# Development of a Charge-Resistant Embedding Media for High-Performance Serial Block Face Imaging of Cells and Tissues

**DOI:** 10.1101/2025.05.28.656661

**Authors:** Thomas Deerinck, Steven Peltier, Mark Ellisman

## Abstract

Serial block-face scanning electron microscopy (SBEM) enables high-resolution 3D imaging of biological specimens but is often limited by specimen charging due to the use of non-conductive epoxy resins. While heavy metal staining and variable-pressure SEM can reduce charging, these methods compromise resolution or are only partially effective. We report a novel approach using polyethylene glycol (PEG 3350), a water-soluble, non-conductive polymer, as a doping agent to reduce specimen charging without sacrificing imaging quality. Although PEG 3350 alone lacks the mechanical and sectioning properties required for SBEM, we found it can be dissolved in standard Durcupan epoxy resin to improve charge resistance while maintaining physical integrity. Resins doped with 10% PEG 3350 demonstrated a substantial reduction in charging at 1.4–1.8 keV in charge-prone samples such as cultured HeLa cells, lung, and brain tissues, while retaining transparency and sectionability. Unlike other methods, the doped resin performs in high vacuum without compromising signal-to-noise ratio or spatial resolution. Notably, the PEG-doped resin reduced charging at voltages above the typical threshold for undoped resins, which show artifacts above 1.0 keV. Though some charging persisted in lung tissue at 2.0 keV, near-complete mitigation was achieved at slightly lower voltages. Our findings suggest that PEG-doped resins provide a promising route to minimize specimen charging in SBEM, and future studies optimizing PEG molecular weight and concentration could yield a universally charge-resistant embedding medium compatible with high-resolution imaging across diverse sample types.

## Introduction

Serial block-face scanning electron microscopy (SBEM) is becoming the method of choice for obtaining large-scale 3-D imaging data of biological specimens at nanometer-scale resolution. The instrument consists of an ultramicrotome fitted within a backscatter-detector equipped SEM. In an automated process, the ultramicrotome removes an ultra-thin slice of the sample surface with an oscillating diamond knife and the region of interest on the block face is imaged using backscattered electrons. This sequence is repeated hundreds or thousands of times until the desired volume of the sample is traversed. This method enables the reconstruction of thousands of cubic microns of tissue at a level of resolution far better than that obtainable by light microscopy and is rapidly becoming the method of choice for many biological research applications, such as neuroscience, cancer research, and general cell biology.

One of the principle limitations of this method is that specimens must be embedded in non-conductive epoxy resin prior to imaging, and this can lead to substantial specimen charging. Typically, cells and tissues are intensely heavy-metal stained in order to improve backscattered electron yield at low accelerating voltages and to reduce specimen charging (Deerinck et al., 2010). However, not all specimen charging can be eliminated in charge-probe samples such as those containing large open spaces (i.e. lung, kidney and liver tissue) or with low lipid content (muscle, bone, plant, etc.). Most commonly, variable-pressure SEM (VP-SEM) is used for many specimens to minimize charging, but at a significant loss of signal-to-noise and resolution owing to electron-gas interactions (Titze and Denk, 2016).

We recently developed an approach to mitigate specimen charging using focal charge compensation (FCC) by injection of nitrogen gas over the block face surface, neutralizing charging in a man similar to VP-SEM (Figure 1), but without the substantial loss of signal-to-noise and spatial resolution because the overall chamber pressure is at high vacuum (Deerinck et al., 2017). While this is a powerful tool to minimize charging at high vacuum, not all SBEM systems are compatible with this technology. The ultimate goal would be to have an embedding matrix that is intrinsically charge-resistant, and a few reports have been published on encasing biological samples in either silver colloid before resin embedding (Wanner et al., 2016) or in a resin containing carbon black (Nguyen et al., 2016) to reduce charging. However, both of these approaches have important drawbacks, including making the specimen optically opaque, and since neither can penetrate into cells and tissues, they are of only modestly effective.

**Figure 1.**
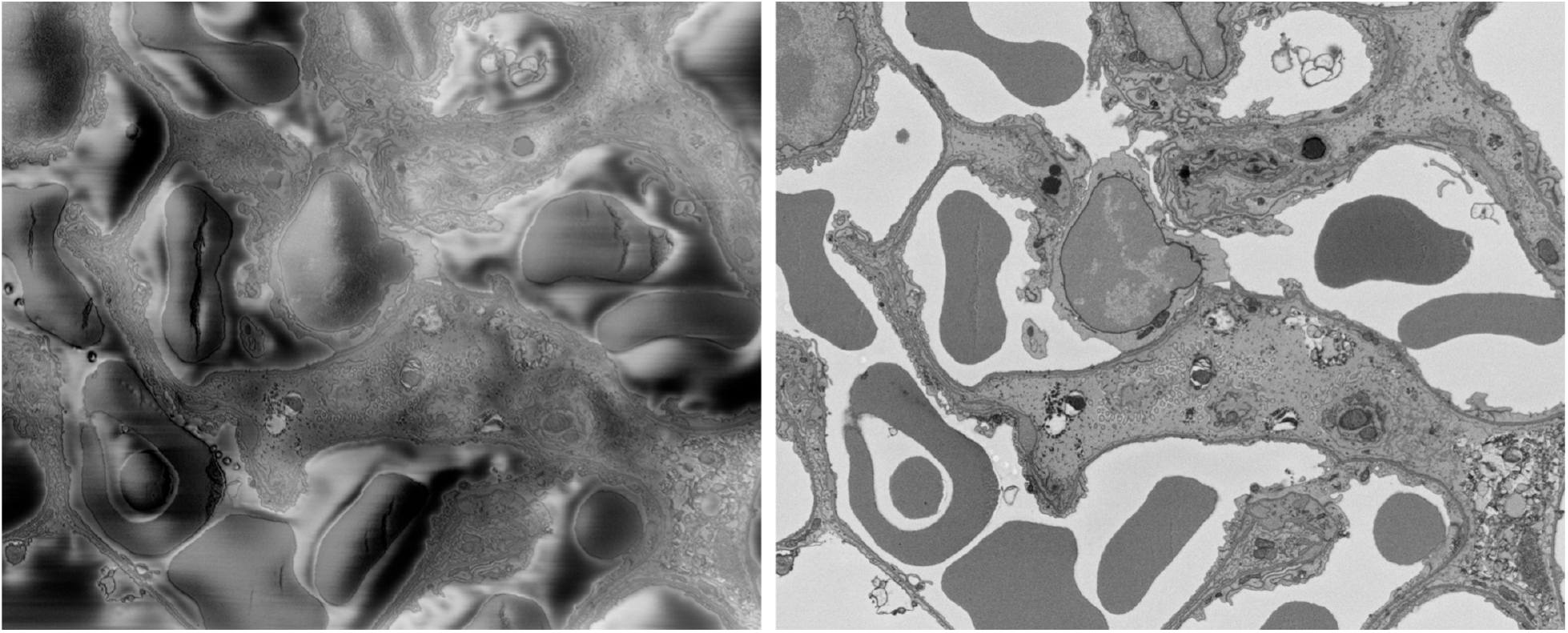
SEM block face images of lung tissue. Left panel: Excessive specimen charging is noted (black streaks) when imaged at high vacuum at 2.5 keV. Right panel: Using FCC, all specimen charging is eliminated. Images from Deerinck et al., 2017.

## Results

Specimen charging in SBEM usually occurs when electrons build up on the specimen surface and electrostatically interfere with the imaging electrons. We recently and inadvertently found that the compound polyethylene glycol M.W. 3350 (PEG 3350) does not exhibit charging at the voltages routinely used for SBEM (1-3keV). PEG 3350 is a water-soluble compound that is used in a wide variety of industrial applications, including the pharmaceutical industry and is the main ingredient in MiraLAX, a common colonoscopy prep solution. It has even been used as an embedding matrix for electron microscopy (Wolosewick 1980), albeit not for SBEM. While PEG 3350 is not conductive, it has the anomalous property of not charging at 1-3 keV, most likely due to the ejection of secondary electrons from the sample elicited by the primary electron beam in great enough number to balance the overall charge, referred to as charge balance (Tang and Joy, 2003). There are other non-conductive materials that have a similar behavior, such as magnesium oxide (Figure 2).

**Figure 2.**
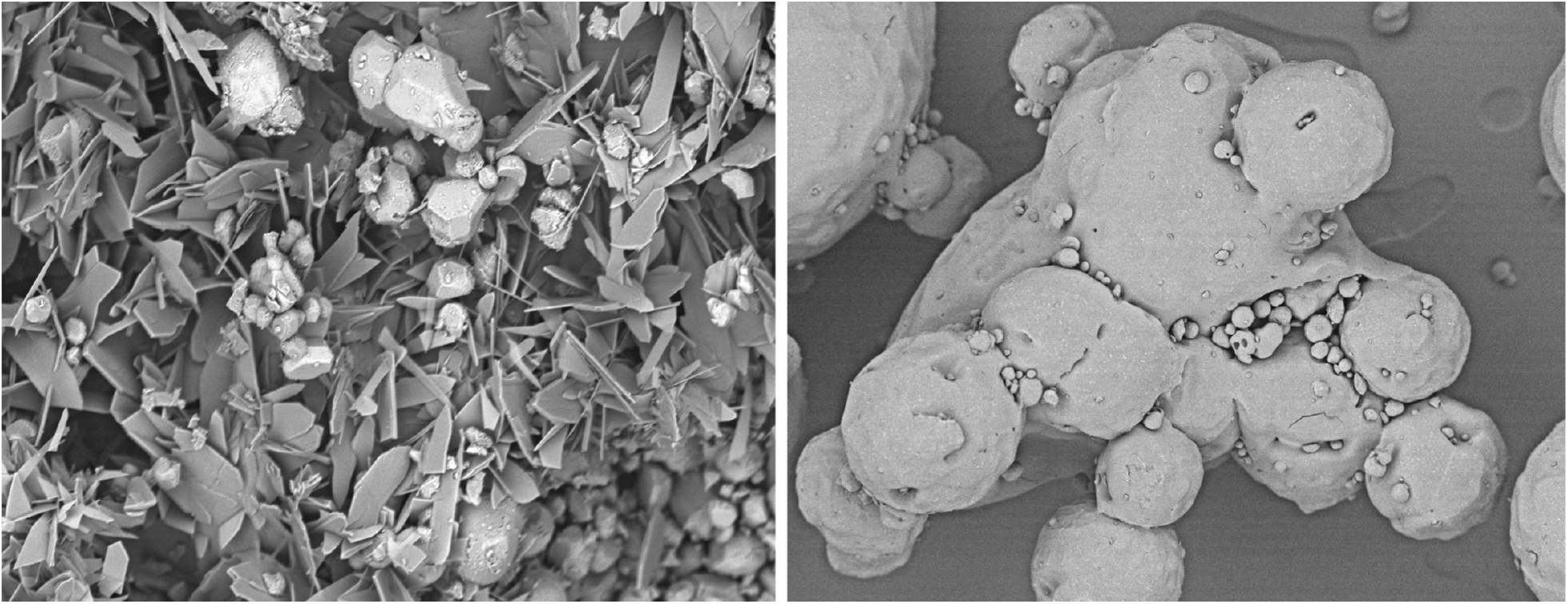
SEM images of uncoated magnesium oxide (left panel) and PEG 3350 (right panel) taken at 3.0 kev accelerating voltage and high vacuum. No specimen charging is observed though neither are conductive materials. Zeiss Merlin FE-SEM.

We have embedded tissues in PEG 3350 and by SBEM found only minimal specimen charging in otherwise extremely charge-prone samples (i.e. lung tissue, Figure 3). However, PEG 3350 itself does not perform as well as the standard epoxy embedding media commonly used with SBEM (Durcupan ACM resin, Fluka), both in terms of beam tolerance and sectionability. We surmised that it might be possible to dope the standard epoxy resin with various amounts of PEG 3350 to raise its charge-balance point and improve its charge-resistance. The goal would be to have a resin with all the positive attributes of epoxy resin (transparent, sections well, beam tolerant, penetrates uniformly into cells and tissues), but is resistant to charging.

**Figure 3.**
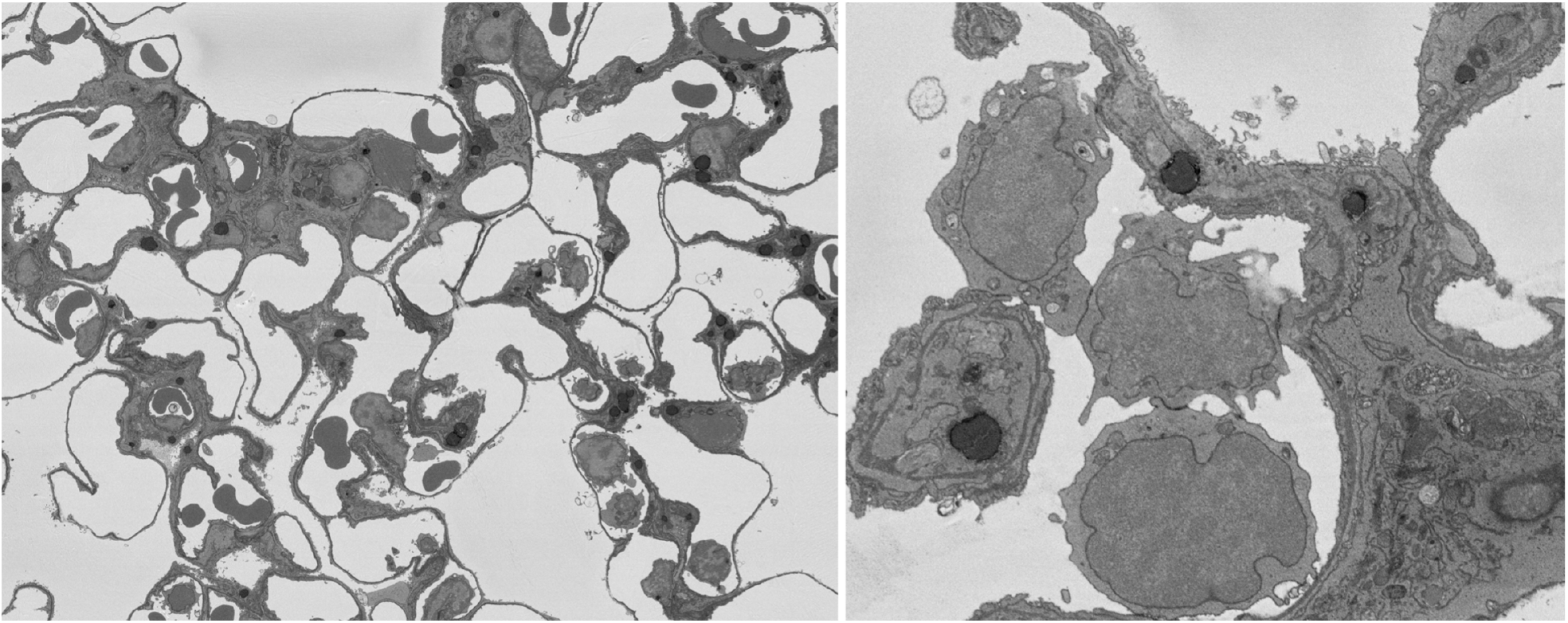
Block face SEM images of lung tissue embedded in 100% PEG and imaged at 2.5 keV. No charging is visible, even in areas of bare PEG. Zeiss Merlin FE-SEM.

PEG 3350 is a solid at room temperature and is soluble in all percentages in epoxy resin, so we began by testing 2, 5, 10, 15 and 20% (w/w) PEG in Durcupan resin. After polymerization, bulk blocks were tested for mechanical strength and sectionability of the resin (Figure 4), and then were placed in the SEM and imaged at a variety of accelerating voltages (1-5 keV) at high vacuum. When compared to undoped resin that shows significant charging at all voltages above 1.0 keV, with the 2% doped resin only modest charging was observed and at 20% almost no charging was observed up to 2.0 keV. However, the 20% resin had the disadvantage of being soft in consistency and therefore reduced in its sectionability. It also became slightly opaque if allowed to cool to room temperature, complicating the infiltration and embedding protocol.

**Figure 4.**
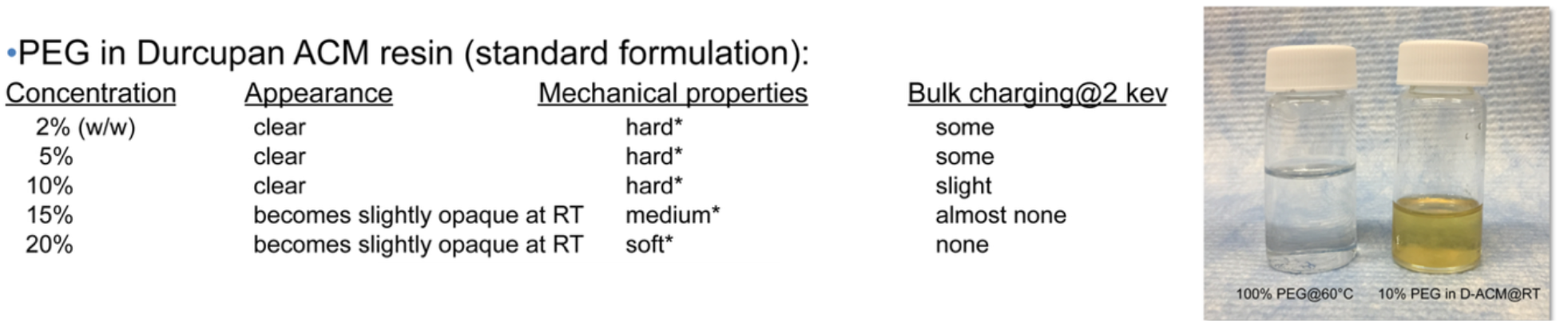
Physical and mechanical properties of various concentrations of PEG 3350 in Durcupan epoxy resin. 10% doped resin remained clear at room temperature.

We decided to test 10% PEG in resin using fixed and embedded cultured HeLa cells grown in a monolayer: a very typical type of sample and charge-prone such that it is only imageable by VP-SEM or FCC. In direct comparison using parallel-processed samples and identical imaging parameters (Zeiss Merlin FE-SEM, ~80pA probe current, HR mode, 8kx12k raster with 1 µsec dwell time, 6.0mm WD, 1730x magnification, chamber vacuum ~1×10^−6^mbar), we found a significant improvement in charge-resistance in the doped resin.

While moderate charging was noted at 2.0 keV, almost no charging was observed at 1.4 and 1.5keV in the doped resin (Figure 5 and 6). The undoped resin in contrast showed significant charging at any voltage above 1.0 keV. The sectionability of the doped resin was comparable to undoped resin when polymerized at 90°C for 24 hours and remained glassy-clear and otherwise indistinguishable from undoped resin.

**Figure 5.**
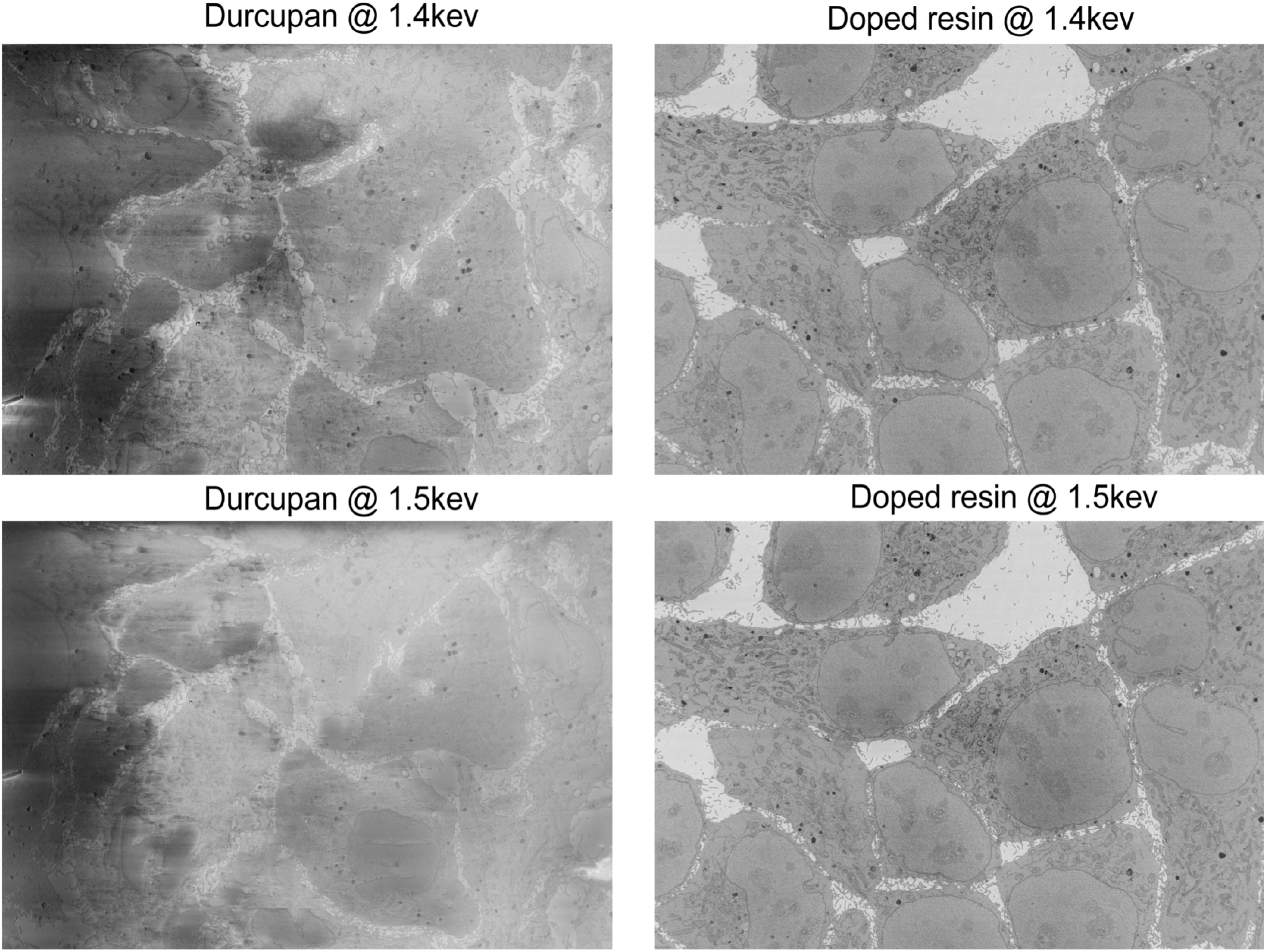
Block face image of cutured HeLa cells with either undoped (left panels), or 10% doped (right panels) resin at two different accelerating voltages at high vacuum.

**Figure 6.**
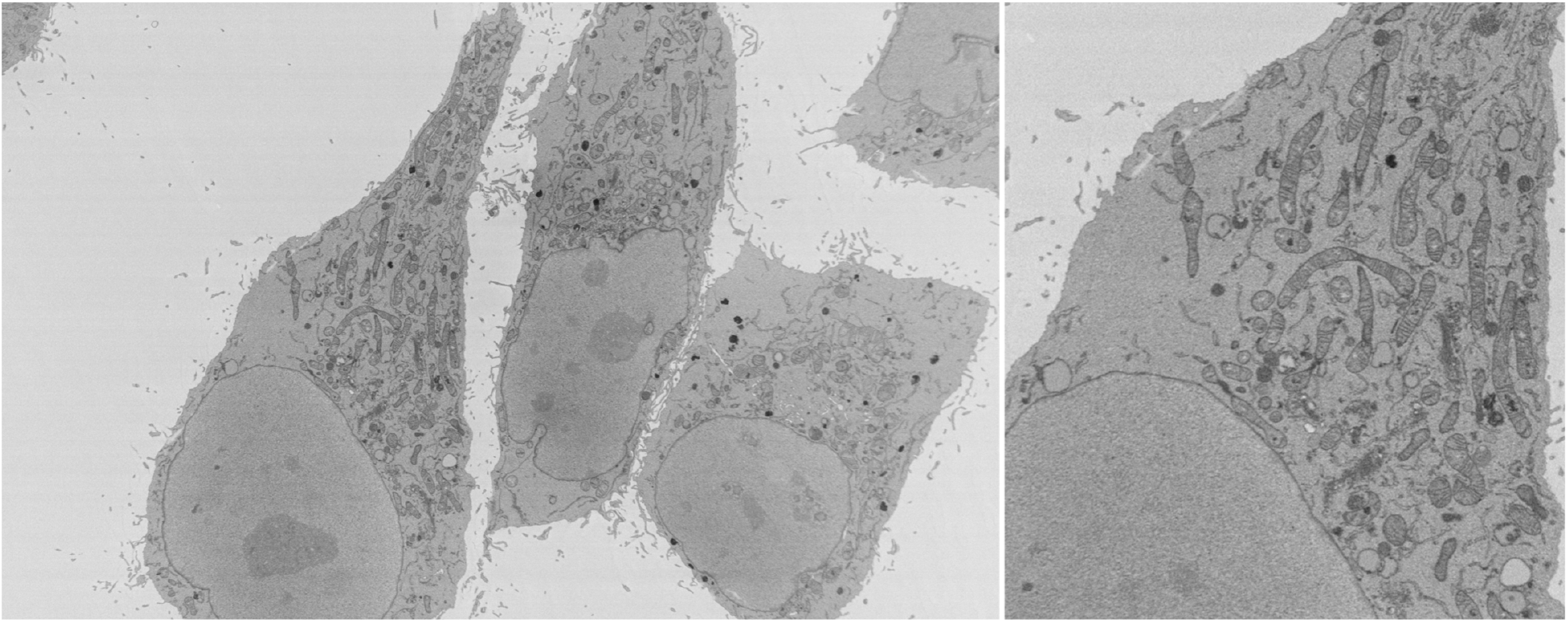
Block face image of cutured HeLa cells using 10% doped resin at 1.4 kev accelerating voltage and high vacuum. No specimen charging is observed.

We next decided to test this resin on both lung and brain tissue. Specimens that were conventionally fixed and prepared using the OTO procedure and *en bloc* uranyl acetate treatment (the same used for the HeLa cells) were embedded in the 10% doped resin and cured at 90°C for 24 hours. We found that at 2.0 keV some specimen charging was noted in the lung tissue but at

1.8 keV essentially no charging was observed (Figure 7). This is in contrast to undoped resin, which showed charging artifacts above 1.0 keV. Brain tissue is more difficult to assess since it can be metal impregnated to such a degree that charging is only visible in blood vessels, capillaries and in some cell nuclei. In parallel with processing lung tissue, we also processed mouse cerebellar tissue (Figure 8).

**Figure 7.**
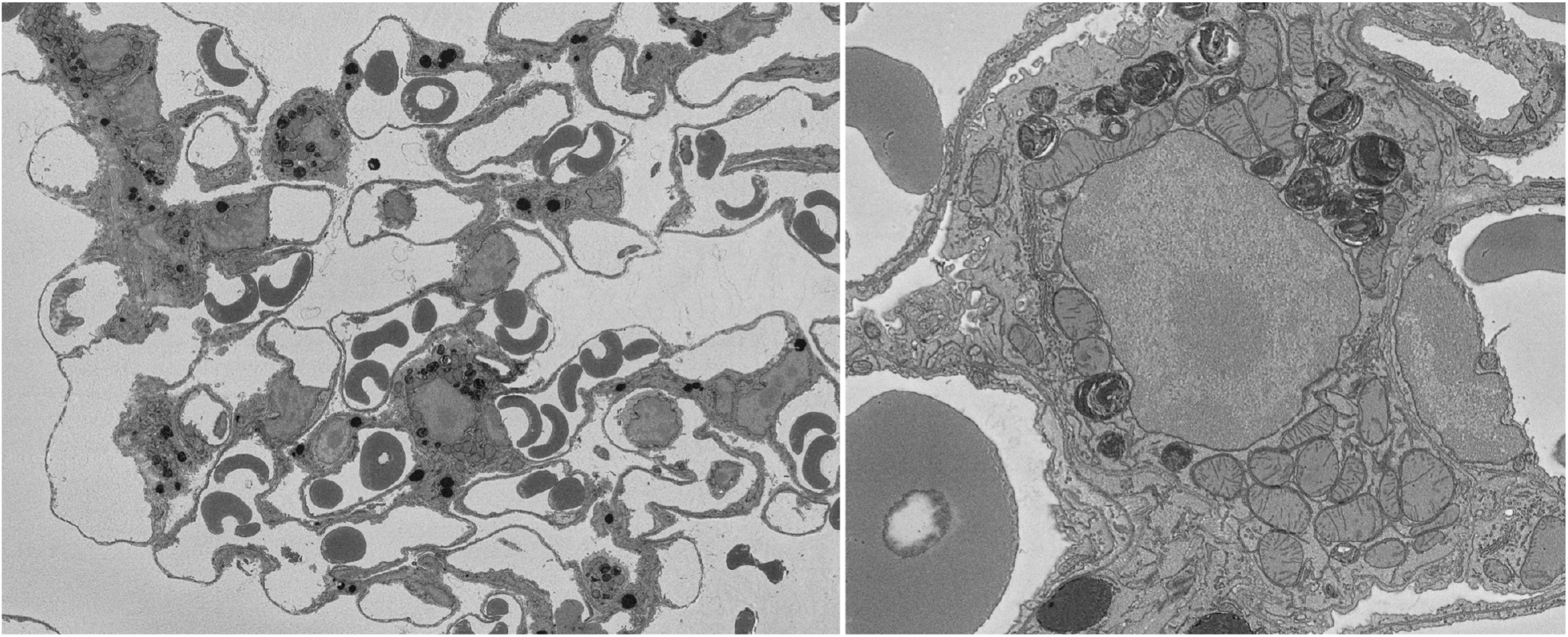
Block face images of lung tissue embedded in 10% doped resin and imaged at 1.8 keV and at high vacuum. No charging is evident. Zeiss Merlin FE-SEM.

**Figure 8.**
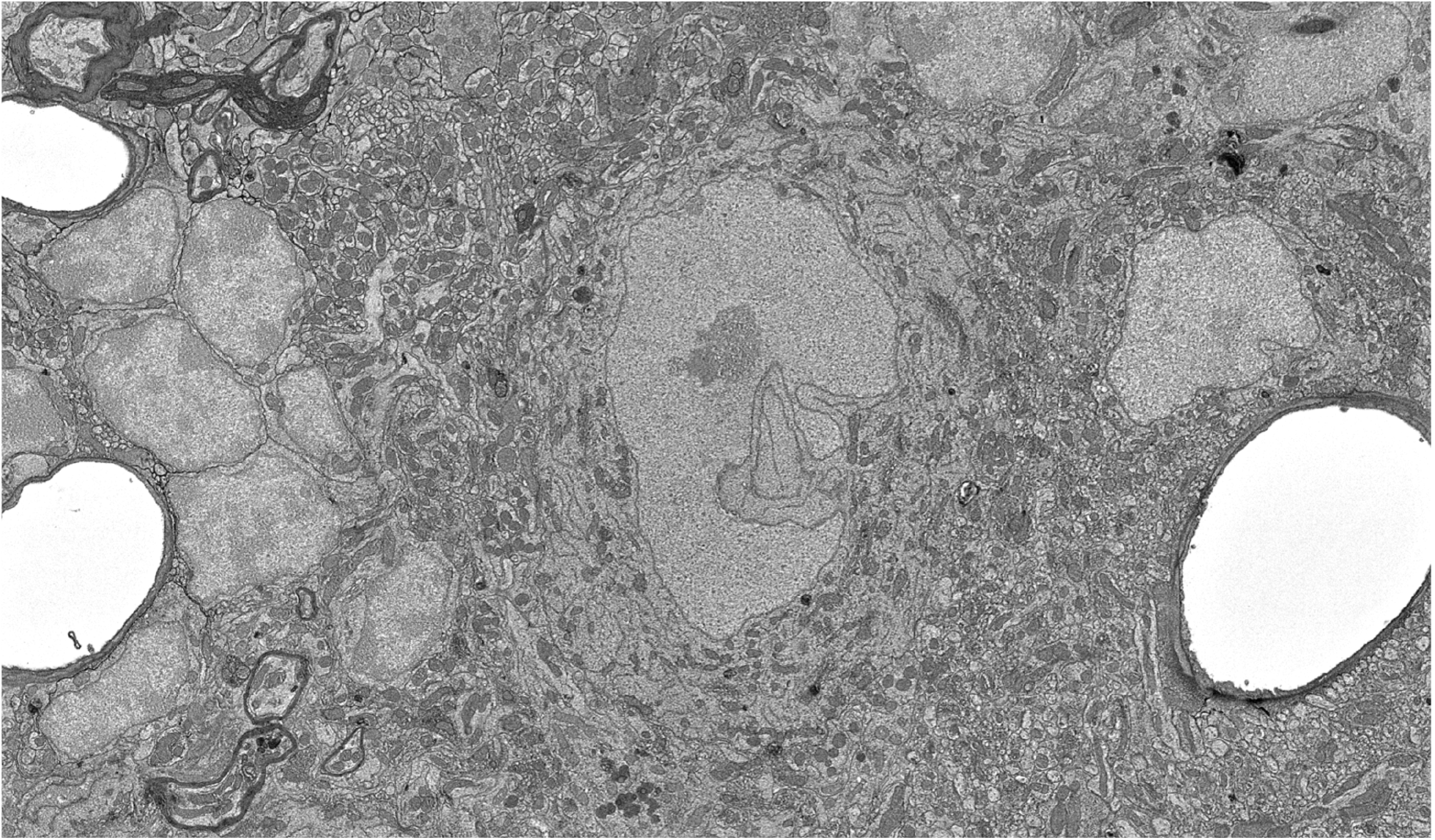
Block face images of cerebellum embedded in 10% doped resin and imaged at 2.0 keV and at high vacuum. No charging is evident in capillaries. Zeiss Merlin FE-SEM.

## Conclusions

Specimen charging is an important limitation in SBEM. While several solutions have been put forward (Nguyen et al., 2018), as yet no universally applicable approach has been found. We believe that the development of a charge-resistant embedding media will play an important role in eliminating this problem. We have found that the addition of PEG 3350 into epoxy resin reduces charging in the accelerating voltage range most commonly used for SBEM. PEG is commercially available in a wide range of average molecular weights, from 400 to over 100,000. We chose PEG 3350 as a starting point since in is a liquid at 56°C, making it compatible with normal infiltration and embedding protocols. While we did not reach our ultimate goal of charge-resistance up to 2.0 keV with all samples, we did see a substatial improvement in the reduction of specimen charging and we believe future work on finding the optimal PEG concentration and molecular weight(s) incorporated into the resin will ultimately result in a doped resin meeting this criteria. We will also need to test PEG in combination with other epoxy resin formulations.

## Acknowledgements

This research was supported by the National Institutes of Health (NIH) BRAIN Initiative, grant number (U24NS120055) to MHE.

## Notes

### Competing Interest Statement

The authors have declared no competing interest.

